# Development and Characterization of Patient-Derived Xenografts from Non-Small Cell Lung Cancer Brain Metastases

**DOI:** 10.1101/2020.06.02.130062

**Authors:** Andrew M. Baschnagel, Saakshi Kaushik, Arda Durmaz, Steve Goldstein, Irene M. Ong, Lindsey Abel, Paul A. Clark, Ticiana Leal, Darya Buehler, Gopal Iyer, Jacob G. Scott, Randall J. Kimple

**Affiliations:** Department of Human Oncology, School of Medicine and Public Health, University of Wisconsin, Madison, WI, USA; University of Wisconsin Carbone Cancer Center, School of Medicine and Public Health, University of Wisconsin, Madison, WI, USA; Case Western Reserve University, Cleveland, OH 44195, USA; Department of Biostatistics and Medical Informatics, University of Wisconsin School of Medicine and Public Health, University of Wisconsin, Madison, WI, USA; Department of Obstetrics and Gynecology, University of Wisconsin School of Medicine and Public Health, University of Wisconsin, Madison, WI, USA; Department of Medicine, Division of Hematology/Oncology, School of Medicine and Public Health, University of Wisconsin, Madison, WI, USA; Department of Pathology and Laboratory Medicine, University of Wisconsin School of Medicine and Public Health, University of Wisconsin, Madison, WI, USA; Department of Radiation Oncology, Taussig Cancer Institute, Cleveland Clinic, 10201 Carnegie Ave, Cleveland, OH 44195, USA

**Keywords:** non-small cell lung cancer, brain metastasis, patient-derived xenografts

## Abstract

**Introduction:** The purpose of this study was to establish and characterize a direct-from patient-derived xenograft (PDX) model of non-small cell lung cancer (NSCLC) brain metastases.

**Methods:** Surgically obtained tissue was implanted subcutaneously and as orthotopic intracranial implants into immunodeficient mice. Histology and DNA loci were compared between original tumor and subsequent PDX passages. Tumors underwent RNA and DNA sequencing and relevant therapeutic targets were identified. Tumor growth rates were assessed following treatment with radiation, MEK inhibitor selumetinib, or MET inhibitor savolitinib. Cell lines were established.

**Results:** Nine NSCLC brain metastases PDXs were established. Morphologically, strong retention of cytoarchitectural features was observed between original patient tumor and subcutaneous and intracranial tumors. Short tandem repeat analysis demonstrated strong concordance between patient tumors and subsequent PDX passages. Transcriptome and mutation analysis revealed high correlation between matched patient and PDX samples. Significant growth inhibition occurred with radiation, with selumetinib in tumors harboring *KRAS* G12C mutations and with savolitinib in a tumor with *MET* exon 14 skipping mutation. The combination of radiation and savolitinib resulted in significant tumor growth delay compared to radiation or savolitinib alone our MET exon 14 skipping mutation PDX. Early passage cell strains showed high consistency between patient and PDX tumors.

**Conclusion:** We have established a robust human xenograft model system for investigating NSCLC brain metastases. These PDXs and cell lines show strong phenotypic and molecular correlation with the original patient tumors and provide a valuable resource for testing preclinical therapeutics.

## Introduction

Brain metastases from non-small cell lung cancer (NSCLC) are unfortunately a common event that occur in more than a quarter of patients with stage IV disease at diagnosis^1^, and up to 30% of patients developing central nervous system (CNS) progression during the course of their disease.^1–3^ Once diagnosed, the prognosis remains poor, with a median survival of 9-15 months.^4^ Treatment options are evolving as patients treated with systemic therapies have better control of their extracranial disease and live longer, which allows time for brain metastases to develop.

Standard treatments for brain metastases are surgical resection and stereotactic radiosurgery for limited lesions or whole brain irradiation for multiple metastases.^5^ Whole brain irradiation has significant toxicity^5^ and cytotoxic chemotherapy has a limited role due to its inability to cross the blood-brain barrier.^5,6^ A proportion of patients with brain metastases harboring alterations in *EGFR* or *ALK* can achieve intracranial control with small molecule tyrosine kinase inhibitors^7,8^; however most patients with NSCLC do not harbor these genomic alterations. In a phase II study, patients with NSCLC and brain metastases with PD-L1 expression of at least 1%^9^ benefited from PD-1 blockade, with CNS responses (30%) similar to extracranial responses, indicating that immunotherapy can results in prolonged survival in a subset of patients. However, these response rates are far below those seen following standard of care radiotherapy. Overall, there is a need to develop therapies that have better intracranial efficacy with less toxicity.

One barrier to improving the therapeutic approach to brain metastases is the lack of robust preclinical brain metastasis models. To our knowledge, there are currently no commercial NSCLC brain metastasis cell lines available and *in vivo* brain metastasis models are not widely accessible. Patient-derived xenografts (PDXs) are a well-established model to study cancers in the preclinical setting.^10^ PDXs are established by obtaining tumor samples directly from patients and subsequently implanting and passaging these tumors in immunodeficient mice.^10^ PDXs can be grown in the flank or hind leg of mice or as orthotopic brain implants. For brain tumors, orthotopic intracranial implants allow cancer cells to more closely resemble their original patient tumors both phenotypically and genotypically given the influence of the brain microenvironment.^11^

Here, we report on the development and characterization of a cohort of NSCLC brain metastasis PDX models, including both flank and orthotopic intracranial xenografts as well as two NSCLC brain metastasis-derived cell lines. We demonstrate the utility of these PDXs by assessing their response to radiation and selumetinib in *KRAS* mutated xenografts and savolitinib in a *MET* exon 14 mutated xenograft.

## Materials and Methods

### Patient Selection

Patients with newly diagnosed or recurrent NSCLC brain metastases, undergoing surgical resection, were consented to participate in this IRB-approved protocol.

### Mice

Immunodeficient NOD SCID gamma (NSG) (Envigo, Indianapolis) and Hsd:athymic Nude-Foxn1nu (Envigo, Indianapolis) mice were used for PDX development. All mice were kept in the Association for Assessment and Accreditation of Laboratory Animal Care approved Wisconsin Institute for Medical Research Animal Care Facility, and studies were carried out in accordance with an approved animal protocol.

### DNA Sequencing

After expert pathologist assessment, solid tumor genomic DNA was isolated from formalin-fixed, paraffin-embedded tumor samples. DNA libraries were prepared using the KAPA Hyper Prep Kit, hybridized to the xT probe set, and amplified with the KAPA HiFi HotStart ReadyMix. The amplified target-captured DNA tumor libraries were sequenced to an average target depth of 500x on an Illumina HiSeq 4000 using the Tempus xT (Tempus labs Chicago, IL, USA) gene panel. The panel analyzes single nucleotide variants (SNVs), indels, and copy number variants in 596 genes and genomic rearrangements in 21 genes with an average coverage of 500x.^12^ Variants were called using Freebayes (version 1.0.2). Clinically relevant mutations were identified using the Cancer Genome Interpreter^13^ and ClinVar^14^.

### Establishment of Xenografts

Patient tissue collected during surgery was added to a 1.5ml tube with media (Dulbecco’s Modified Eagle Medium with 10% fetal bovine serum, 1% penicillin/streptomycin, and 2.5 μg/mL amphotericin B) and minced with sharp, sterile scissors into a slurry. Matrigel (#354230, BD Biosciences, Inc) was added to the media in a 1:1 ratio and injected subcutaneously into NSG mice with an 18 gauge needle. Subsequent passages were made in a similar fashion into either NSG or athymic nude mice. Cryopreservation of xenografts was accomplished by mixing minced tumor pieces with complete transport media supplemented with 10% dimethylsulfoxide (DMSO) as previously described.^15–18^ Tumors were frozen in controlled rate freezers to –80°C overnight and transferred to liquid nitrogen for long-term storage. To thaw tumors, aliquots were warmed to 37°C in a heated water bath and tumor tissue was washed twice in transport complete media (without DMSO) and immediately implanted into mice.

### Establishment of Cell Lines

Tumors taken from early PDX passages were dissociated into single cell suspensions and seeded in T-25 flasks in media (Dulbecco’s Modified Eagle Medium with 10% fetal bovine serum, 1% penicillin/streptomycin, and 2.5 μg/mL amphotericin B) and incubated at 37°C in a humidified atmosphere of 5% CO2. Media was changed regularly, and cells were passaged once they reached 70% confluence. Mycoplasma testing and short tandem repeat profiling was performed during passaging.

### Short Tandem Repeat (STR) Profiling

DNA typing by multiplex polymerase chain reaction (PCR) of short repetitive elements of highly polymorphic markers was performed to assess genetic stability of xenografts and cell lines. STR profiling and analysis was performed using the PowerPlex® 16 HS System (Promega, Madison WI).

### Orthotopic Intracranial Implants

Tissue was enzymatically dissociated to single cells and 2×10^5^ cells were suspended in 5 μl of phosphate-buffered saline. Using a Hamilton syringe, the cells were stereotactically injected under sterile conditions into the right striatum of anesthetized NOD SCID mice at 1 μL/min at the following coordinates referenced from bregma: 0 mm antero-posterior, +2.5 mm medio-lateral, and −3.5 mm dorso-ventral. The scalp was closed with 6-0 polypropylene suture. Mice were placed on a heating pad in sterile cages and allowed to awaken from anesthesia. One week after injection animals were placed onto a small-animal MRI scanner (4.7-T horizontal bore imaging/spectroscopy system; Varian, Palo Alto, CA), and T1- and T2-weighted images were obtained. As per animal protocol, once neurological symptoms were observed, mice were immediately euthanized by perfusion fixation with 10% paraformaldehyde and moved to 70% ethanol solution after 48 hours. Brains were then excised, formalin fixed, embedded in paraffin, and processed for histology.

### Immunohistochemistry

At each passage, tumor was fixed in 10% neutral-buffered formalin and embedded in paraffin blocks. Five-μm sections of PDX and matching patient tumors were stained with hematoxylin and eosin (H&E) and imaged on an Olympus BX51 microscope (Olympus America, Inc). TTF-1 (clone SPT24, BioCare, 1:100) staining was performed with a BenchMark Ultra IHC/ISH System (Roche Diagnostics, Rotkreuz, Switzerland); according to manufactures established protocol. Additional Immunohistochemistry (IHC) details and antibodies used can be found in the supplemental materials.

### RNA Sequencing

Total RNA was isolated by using the QIAGEN AllPrep Kit. RNA sequencing libraries were prepared using the KAPA Stranded RNA-Seq Kit with RiboErase (HMR; KAPA Biosystems). RNA was fragmented to a size of 100–200 nucleotides and amplified for 11 cycles. Final libraries were quantified on a high-sensitivity bioanalyzer chip and sequenced on the NextSeq 550 (Illumina).

### Gene Expression and Pathway Analysis

RNA-Seq data were aligned to human reference sequence GRCh38 using RSEM^19^, and expression quantification per gene was computed. Raw read counts were normalized by the trimmed mean of M-values method^20^. Differential gene expression (DE) analysis was performed using a likelihood ratio test with the R package edgeR^21,22^. The p-values were adjusted to control for false discoveries using the Benjamini-Hochberg method^23^. Principal component analysis (PCA) was performed on log2 transformed count per million values. The gene lists were sorted as increasing or decreasing. For the increasing rankings, the genes with positive log fold change (logFC) were sorted by q-value in increasing order, followed by genes with negative logFC sorted by q-value in decreasing order. For the decreasing rankings, the increasing list was reversed.

Gene ontology (GO) and gene set enrichment analyses (GSEA) were performed using goseq^24^ and GSEAPreranked^25^, and p-values were adjusted with Benjamini-Hochberg method. The GOseq analysis was performed using the annotation package Bioconductor TxDb.Hsapiens.UCSC.hg38.knownGene: Annotation package for TxDb object(s). (Team BC, Maintainer BP (2019), R package version 3.4.6.). GSEAPreranked was performed using the KEGG subset of the canonical pathways gene sets from MSigDB version 7^26,27^. For identifying annotations overrepresented in the supervised approach, the top 1% (219) of the genes were extracted from the DE increasing rankings; in the unsupervised approach, the top 1% (219 genes) were extracted from the PC2 increasing rankings. For identifying annotations overrepresented in the patient tumors increasing rankings were used and for PDX samples decreasing rankings were used. To mitigate the high false discovery rate with GSEAPreranked^28^, a filtering step as described by Yoon et al.^28^ was implemented. Pearson correlation coefficient of the log2 transformed gene expression values was computed, first using all of the genes for each pair of samples and then excluding the top 1% of genes from the PC2 increasing rankings.

### Mutation calling from RNA-Seq

Bulk RNA-Seq files are pre-processed for quality control using fastp with default parameters^29^. STAR aligner in two-pass mode is used to align the reads to GRCh38 and GRCm38 genomes^30^. Aligned bam files for PDX data are further filtered to remove reads aligning to the host genome with high confidence using R package XenofilteR^31^. Following read splitting and duplicate removal, we have utilized Strelka2 in rna mode to call variants with default parameters^32^. Variants passing the default filters and read depth 20 are annotated using Ensembl VEP and filtered to include only high impact variants for downstream analysis.

### Tumor Growth Delay Experiments

Tumor cells (5 × 10^6^) suspended in saline were injected subcutaneously into the hind leg. When tumors grew to a mean volume of 150 mm^3^, mice were randomized into control group, radiation alone group (2 Gy per fractions in 1-2 weeks), selumitinib group (25 mg/kg BID for 2 weeks), savolitinib group (2.5 mg/kg for 2 weeks) or radiation plus savolitinib group (2.5 mg/kg for 2 weeks). Selumitinib and savolitinib were administered by PO gavage. Each experimental and control group contained 10 mice. Tumor volume was measured twice weekly with Vernier calipers and calculated using: 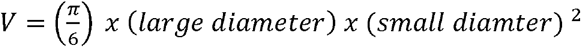. Tumors were followed until the group reached a mean size of 1,500-2,000 mm^3^. Specific tumor growth delay was calculated as the number of days for the mean of the treated tumors to grow to 500 mm^3^ minus the number of days for the mean of the control group to reach the same size. The percentage of tumor growth inhibition (%TGI = 1 – [changes of tumor volume in treatment group/changes of tumor volume in control group] × 100) was used for the evaluation of anti-tumor efficacy. The radiation dose enhancement factor was calculated by dividing the normalized tumor growth delay in mice treated with both savolitinib and radiation by the absolute growth delay in mice treated with radiation alone. Normalized tumor growth was defined as the time in days for tumors to reach 1,000 mm^3^ in mice treated with the combination of savolitinib and radiation minus the time in days in mice treated with savolitinib alone.

### Statistical Analysis

Differences between tumor mean values was determined using a Student’s t-test. Data are presented as the mean ± standard error of the mean (SEM). A probability level of a p value of <0.05 was considered significant. All graphs and analyses were performed and graphed using GraphPad Prism version 8.0 (GraphPad Software, San Diego, CA).

## Results

### Patient Characteristics

Fourteen patients with NSCLC brain metastases were consented for tumor collection and establishment of xenografts. Overall, nine brain metastasis samples have given rise to viable xenograft tumors with a take rate of 64%. Clinical characteristic of the patient and their tumors that successfully formed xenografts are shown in Figure 1. The average age of the donors was 61 (range 38-84). All tumors were adenocarcinomas except one, which was a sarcomatoid carcinoma with an adenocarcinoma (TTF-1 positive) component. The majority (71%) were diagnosed as synchronous brain metastases. Two patients underwent surgical resection for salvage treatment after progression following whole brain irradiation.

**Figure 1:**
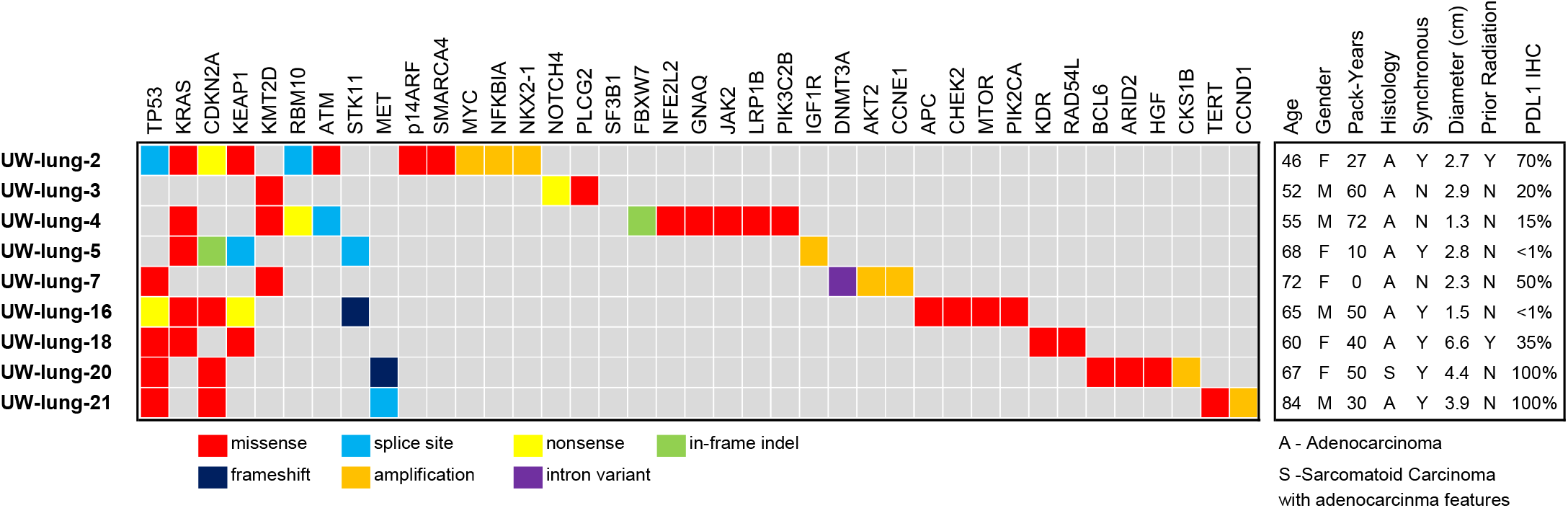
Patient Characteristics and summary of clinically relevant gene mutations from the patient nonsmall cell lung cancer (NSCLC) brain metastasis samples in which patient-derived xenografts (PDXs) were established. Age at diagnosis, gender, smoking pack-years, synchronous brain metastasis, yes or no, prior radiotherapy yes or no, programmed death-ligand 1 (PD-L1) immunohistochemistry (IHC) percent positive.

All nine patient brain metastasis samples underwent amplicon DNA sequencing (Figure 1). The most common genomic alterations were *TP53* mutations in six tumors and *KRAS* mutations in five tumors. Other relevant genomic alterations include *MYC* amplification and loss of *CDKN2A.* One tumor also harbored a *MET* exon 14 splice site mutation. Programmed death-ligand 1 (PD-L1) expression of the patient brain metastasis tumors ranged from <1% to 100% (Figure 1 and supplemental Table 1).

### Establishment and Characterization of Xenografts and Cell Lines

All nine PDX tumors were passaged at least 5 times as subcutaneous xenografts with a mean time between passages of 46.4 days (22-114 days). There were no differences between time of passage from initial tumor implantation to first passage versus fourth to fifth passage (p=0.52). Morphologically, strong retention of cytoarchitectural features was observed between patient tumor and subcutaneous PDX. Figure 2 shows representative H&E images of the patient sample and PDX passage 1 (P1) and passage 5 (P5). TTF-1 expression status was also highly consistent between patient tumor and PDX (Figure 2). Genetic profiles of the PDXs were confirmed with STR analysis (Supplemental Table 2). All PDXs showed 89-100% allelic match to the original patient samples. PDXs continued to retain their STR profiles from passage one to passage five (Supplemental Table 2). Comprehensive tumor growth rate assessment was carried out for five PDXs (Figure 3A). Orthotopic intracranial models were successfully generated from two tumor samples; University of Wisconsin (UW)-lung-2 and UW-lung-16. MRI images shows the formation of brain metastasis and adjacent H&E confirmed tumor which was morphologically similar to the patient and flank tumors (Figure 3B). STR testing confirmed 100% and 92.8% matching of alleles for UW-lung-2 and UW-lung-16 orthotopic tumors, respectively. Cell lines were successfully established from two PDXs, UW-lung-2 and UW-lung-16, following at least 15 *in vitro* passages (Figure 3C). STR testing of subsequent passages confirmed that these lines retained their genetic profile (supplemental table 2).

**Figure 2:**
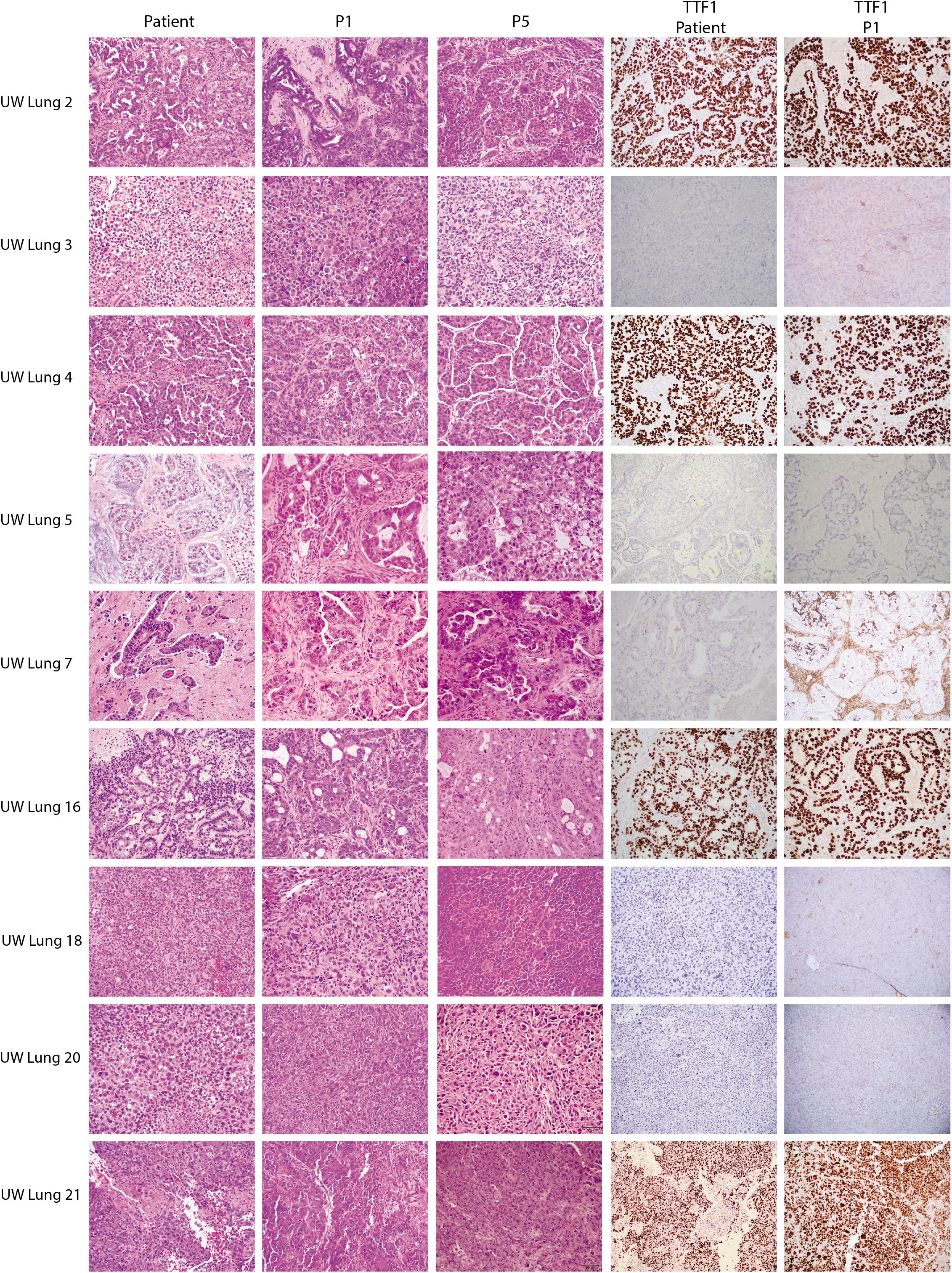
Histopathologic features of nine patient non-small cell lung cancer (NSCLC) brain metastases and corresponding NSCLC brain metastasis patient-derived xenografts (PDXs). Shown are photomicrographs hematoxylin and eosin (H&E) of patient tumor, PDX passage 1 (P1) and passage 5 (P5) and thyroid transcription factor-1 (TTF-1) of patient tumor and passage 1 (P1). Images at 20X.

**Figure 3:**
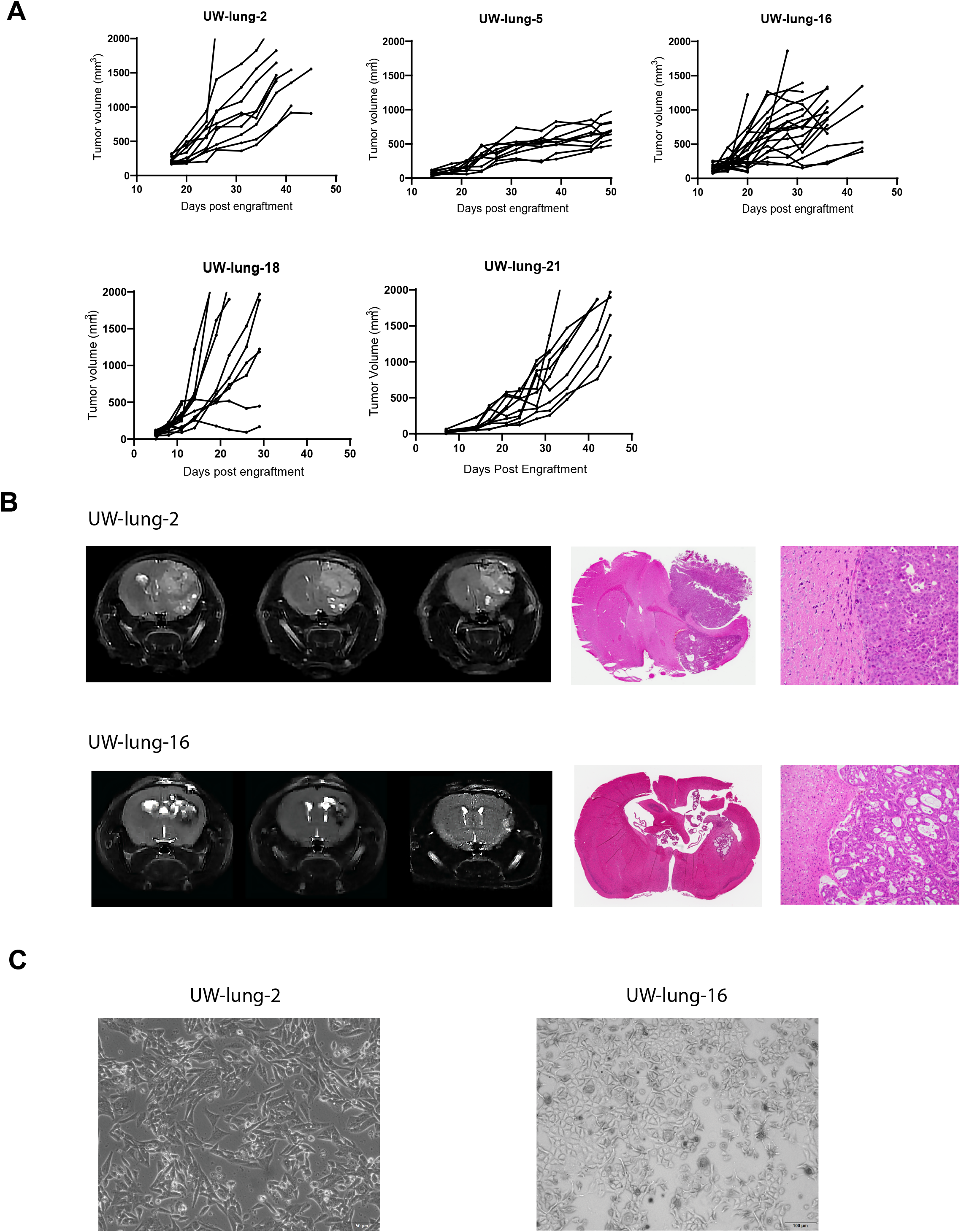
Establishment of subcutaneous flank and orthotopic intracranial patient non-small cell lung cancer (NSCLC) brain metastasis xenografts. **(A)** Subcutaneous flank xenograft growth characteristics of University of Wisconsin (UW)-lung-2, UW-lung-5, UW-lung-16 and UW-lung-18 and UW-lung-21. **(B)** Representative MR images and histology of UW-lung-2 and UW-lung-16 orthotopic intracranial xenografts. **(C)** Pictures of established cell lines UW-lung-2 (passage 20) and UW-lung-16 (passage 20).

### Transcriptome Analysis of patient and PDXs tumors

RNA-Seq analysis was performed on four patient and corresponding flank PDX samples. One PDX (UW-lung-16) was sequenced twice and was used as a control replicate. Mouse-specific reads were filtered in silico from the PDX RNA samples before aligning reads to the human genome. Overall, patient and PDX samples highly correlated with each other (Figure 4A). The average Pearson correlation coefficient for case-matched tumors was 0.88 and ranged from 0.86 to 0.90. Using a supervised analysis comparing matched patient tumors and PDXs, there were 21 Kyoto Encyclopedia of Genes and Genomes (KEGG) pathways that were significantly over represented in patient tumor samples (q < 0.001) and 12 processes that were significantly over represented in the PDX tumors (q < 0.001) (Supplemental Table 3). There were 231 Gene Ontology (GO) annotations overrepresented in patients (q < 0.001) and 185 GO annotations significantly overrepresented in PDX tumors (Supplemental Table 4). The top 10 most significantly overrepresented GO annotations in the patient tumor samples consisted of cellular processes associated with immune regulation while the top pathways overrepresented in the PDXs included those related to cell cycle and DNA replication (Figure 4B).

**Figure 4:**
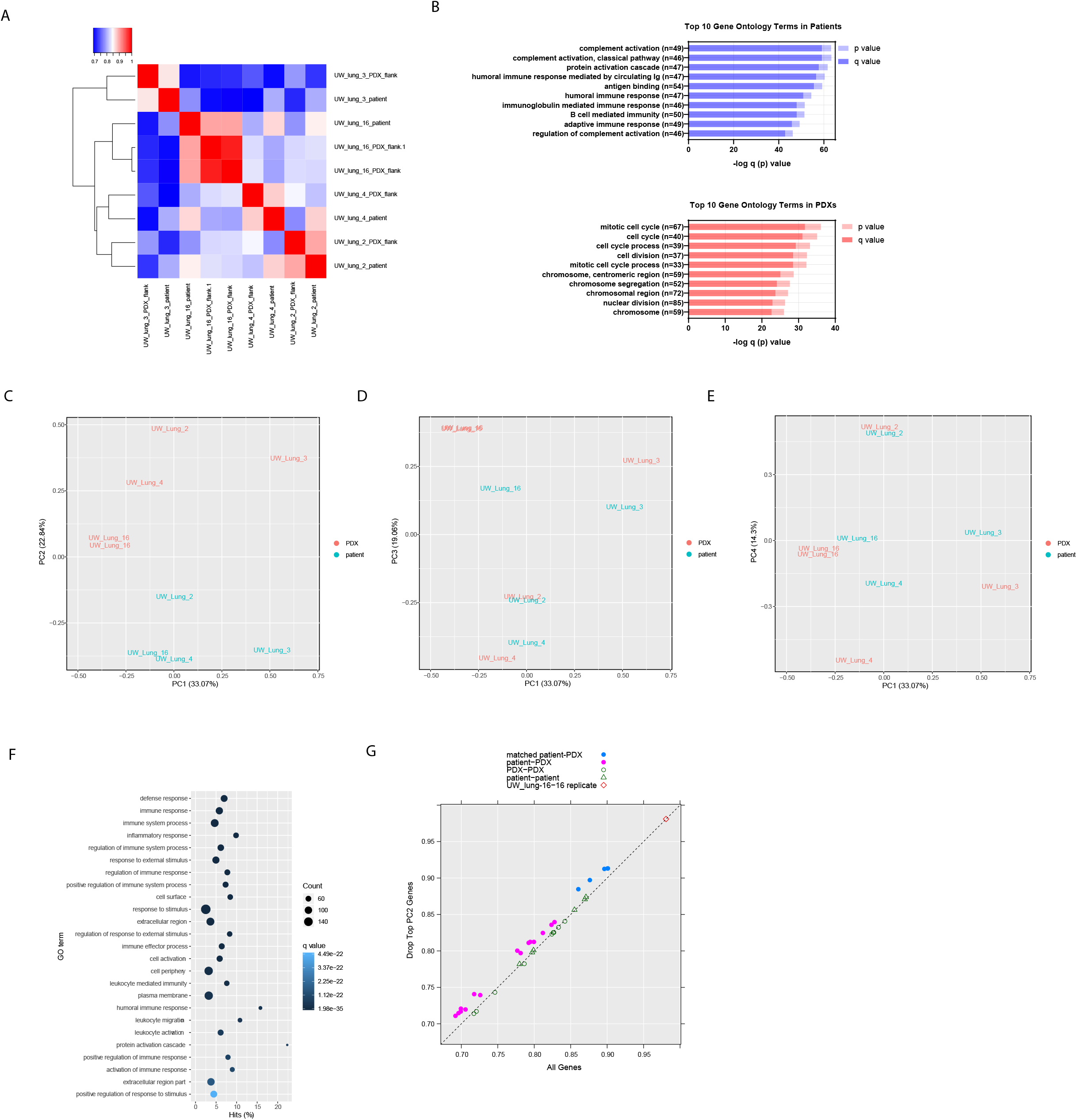
RNAseq shows high correlation between patient and patient-derived xenografts (PDXs) tumor samples. **(A)** Correlation heatmap and hierarchical clustering of patient and PDX tumor samples. **(B)** Top 10 Gene Ontology (GO) terms as summarized by bar charts showing the p value (lighter hue) and false discovery rate (FDR) q value (darker hue). Pathways are labeled along the y-axis; the number of genes annotated within each pathway is indicated in parentheses. **(C-E)** Principal component analysis (PCA) clustering of patient and PDX tumor RNA samples. **(F)** Top GO terms overrepresented in PC2; showing the number of genes (count), the q-value and the percent hits. **(G)** Pearson correlation coefficient of all genes on the x-axis and after removing the top 1% (219 genes) contributing to PC2.

PCA of matched patient and PDX tumor gene expression revealed distinct clustering of PDXs compared to patient tumor samples when viewed through PC1 and PC2 (Figure 4C-E). PC2 appeared to be the main contributor of the differences between patient and PDX samples (Figure 4C). By contrast, viewing the PCA through PC1/PC3 and PC1/PC4 revealed close clustering of matched PDX samples to their corresponding patient sample. (Figure 4D and 4E). On Gene Ontology enrichment analysis, PC2 is highly represented by immune cell processes (Figure 4F). PC1, PC3 and PC4 have little representation of immune processes and appear to represent differences in intrinsic tumor biology (Supplemental Table 4).

Given the differences seen in PC2, we hypothesized that PDXs would associate most similarly to their matched patient tumors compared with unmatched tumors after controlling for the differences in immune cell populations. We performed an analysis removing the top 1% of genes contributing to PC2. This removed 219 genes that were the most prevalent genes associated with immune cell populations (Figure 4F), and a stronger transcriptional resemblance between patient and PDX tumor pairs was seen. The average Pearson correlation coefficient for case-matched tumors and PDXs increased from 0.88 to 0.90 (Figure 4G).

### Patient and PDX Variant Allele Fraction Analysis (VAF)

Variants detected by RNA-Seq for UW-lung-2, UW-lung-4, UW-lung-5 and UW-lung-16 were compared between patient tumors and PDXs grown in the brain and in the flank. High correlations were observed between samples with Pearson correlation coefficients ranging between 0.794 and 0.839 (Figure 5A and B). For UW-lung-2 a comparison between patient brain metastasis and PDX grown in the flank and PDX grown in the brain was also performed. Overall, there were 1,979 alterations detected in the UW-lung-2 patient brain metastasis with 96.5% of these alterations identified in the PDX samples (Figure 5A). There was a higher correlation in VAF distribution (R^2^ = 0.848 and 0.794, respectively) of shared variants, between the patient and the PDX brain metastasis compared to patient and PDX flank tumors. Overall, these findings indicate that the mutation landscape is conserved between patient and PDX samples. In addition, our orthotopic intracranial PDX more closely resembles the originating patient brain metastasis compared to the heterotopic PDX.

**Figure 5:**
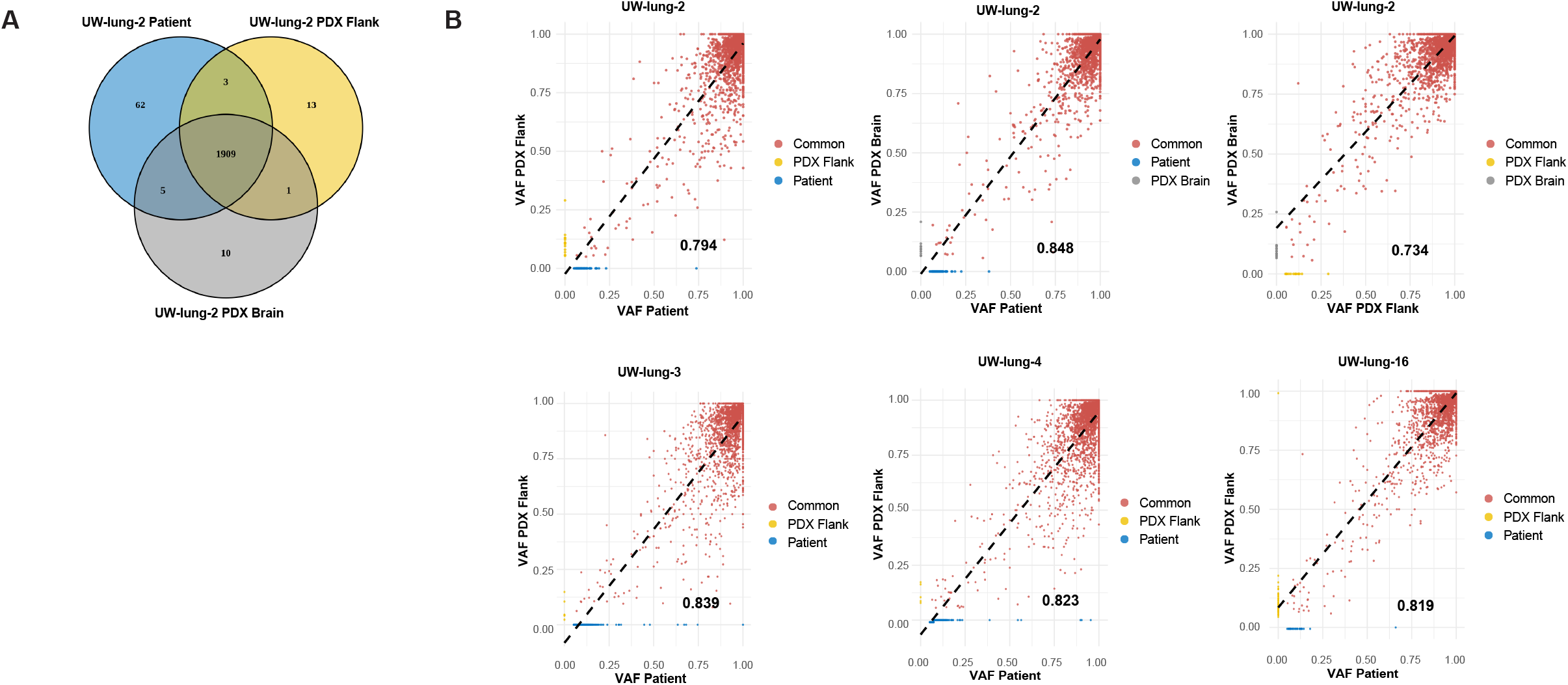
Conservation of mutational landscape. (A) Venn diagram of all variants detected in UW-lung-2 patient brain metastasis, brain PDX and flank PDX. (B) Scatterplots displaying the correlation between patient brain metastasis variant allele frequency (VAF) and flank PDX, patient brain metastasis VAF and brain PDX and flank PDX and brain PDX. The R^2^ value is represented by the black value in the lower right-hand corner of each plot.

### PDX Response to Radiotherapy

Given that radiation is the predominate therapeutic modality for the treatment of brain metastases, five PDXs were tested for response to standard fractionated radiotherapy (Figure 6A). All tumors responded to radiation with tumor growth inhibition values at the end of the experiments of 51% for UW-lung-2, 42% for UW-lung-5, 53% for UW-lung-16, 58% for UW-lung-18 and 92% for UW-lung-21 (P<0.01 for all).

**Figure 6:**
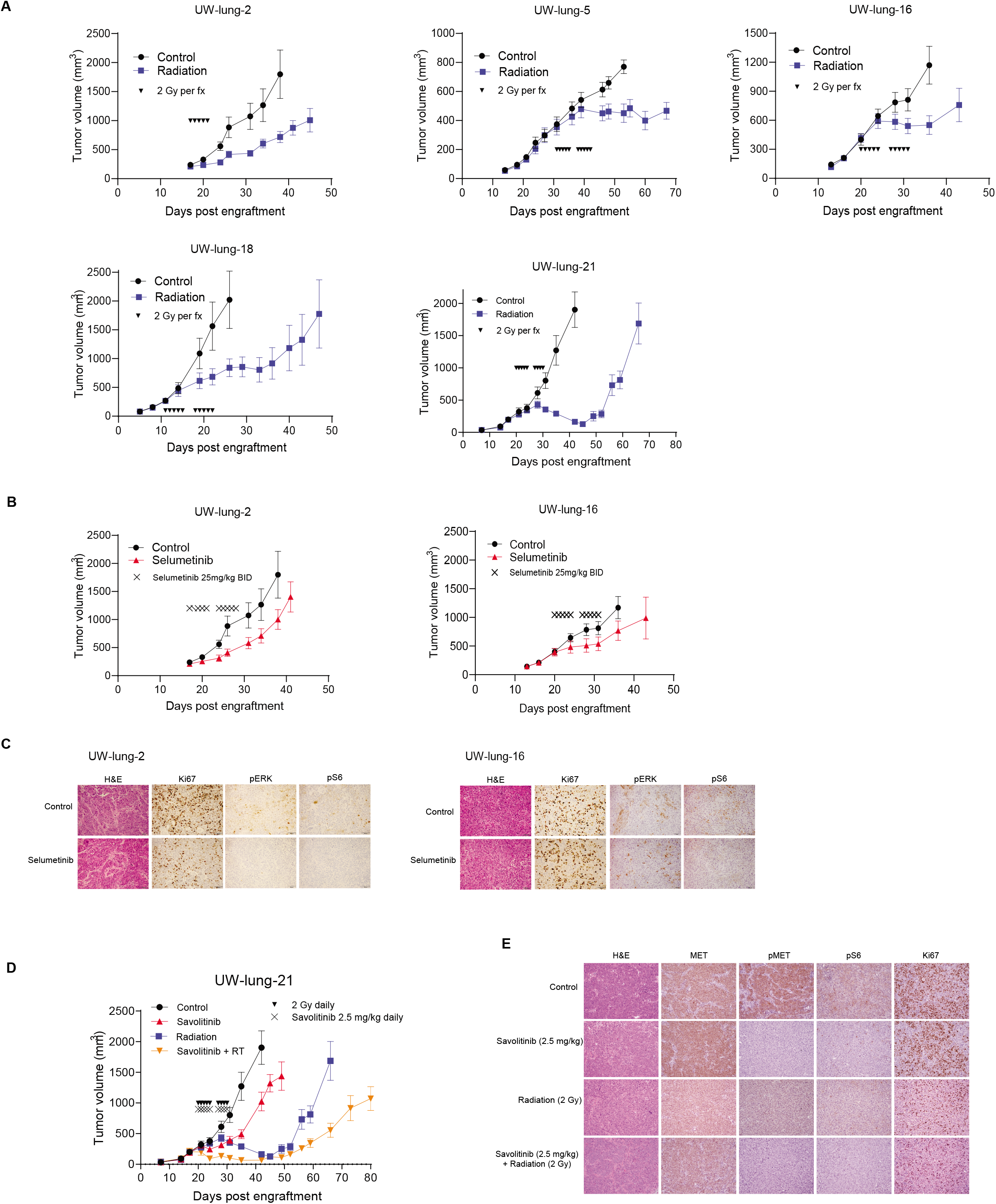
Therapeutic effects of radiation, selumitinib and savolitinib on xenograft growth delay. **(A)** Radiation-induced tumor growth delay curves of UW-lung-2, UW-lung-5, UW-lung-16, UW-lung-18 and UW-lung-21. Radiation was given as 2 Gy per fraction for either 5 days or 10 days as indicated. **(B)** KRAS G12C mutated UW-lung −2 and UW-lung-16 subcutaneous flank xenografts treated with vehicle or selumitinib 25 mg/kg BID for 10 days. **(C)** Immunohistochemistry (IHC) of UW-lung-2 and UW-lung-16 tumor sections showing modulation of KRAS-downstream signaling (p-Erk and p-S6) following a one-time dose of 25 mg/kg BID of selumitnib, collected at 24 hours. **(D)** MET exon 14 mutated UW-lung-21 subcutaneous flank xenografts treated with vehicle, savolitinib 2.5 mg/kg, radiation or radiation plus savolitinib 2.5 mg/kg for 10 days. **(E)** Immunohistochemistry of UW-lung-21 tumor sections showing modulation of MET signaling (MET, p-MET, p-Erk and p-S6) and decrease in Ki-67 following a single dose of 2.5 mg/kg of savolitinib, collected at 24 hours. Points, mean tumor volume in mice after treatment; bars, SEM. The scale bar represents 50 mm and all images are to the same scale.

### PDX Response to MEK Inhibitor Selumetinib

*KRAS* is one of the most commonly mutated genes in lung adenocarcinoma^33^, and in our cohort five PDX tumors harbored *KRAS* mutations. Mutations in *KRAS* leads to activation of the mitogen-activated protein kinase (MAPK) pathway involving MAPK kinase (MEK). As an initial effort to assess the utility of these PDXs for therapeutic response to molecular targeted therapy, we selected two tumors harboring *KRAS* G12C mutations and tested their response to the MEK inhibitor selumetinib. Both tumors responded to selumetinib; however, tumor growth inhibition was only significant in UW-lung-2 (p=0.005) and not significant in the UW-lung 16 (p=0.34) (Figure 6B). IHC analysis revealed a decrease in phospho-MAPK, pS6 and Ki67 with treatment of selumetinib at 24 hours in UW-lung-2. In UW-lung-16, there was minimal change in phospho-MAPK and Ki-67, but a decrease in pS6 (Figure 6C) with selumetinib treatment at 24 hours.

### PDX Response to MET Inhibitor Savolitinib

*MET* exon 14 skipping mutations occur in 3-4% of lung adenocarcinomas and results in the deletion of the intracellular juxtamembrane domain of the receptor, leading to enhanced signaling and tumor proliferation.^33,34^ Small molecule inhibitors have shown therapeutic efficacy in NSCLC harboring MET exon 14 mutations.^34^ We therefore tested the therapeutic response of UW-lung-21, which harbors a MET exon 14 mutation, to the selective MET inhibitor savolitinib. Savolitinib alone significantly inhibited tumor growth in UW-lung-21 with a tumor growth inhibition value of 46% (p<0.01, Figure 6D). We next tested the capacity of savolitinib to augment the response of radiation. Previous studies have shown mixed results when combining radiation with MET inhibitors.^35–37^ The combination of savolitinib and radiation significantly delayed the growth compared to vehicle control, savolitinib or radiation alone (p<0.01; Figure 6D). The mean time for tumors to reach a size of 1,000 mm^3^ was 12.3 days for vehicle-treated mice; 21.6 days for savolitinib-treated mice, 40.5 days for irradiated mice and 54.7 days in mice that received savolitinib plus irradiation (P < 0.001). The absolute growth delay was 9.3 days for savolitinib alone, 28.2 days for irradiation alone and 42.4 days for savolitinib plus irradiation. The radiation dose enhancement factor for savolitinib was 1.2, indicating that savolitinib enhances radiation-induced tumor growth delay. IHC performed on UW-lung-21-treated tumors demonstrated inhibition of phospho-MET and pS6 and decrease in Ki67 with savolitinib treatment (Figure 6E).

## Discussion

Brain metastases from NSCLC frequently occur, remain difficult to treat and if not controlled can lead to significant neurologic compromise and ultimately death.^2^ Current standard approaches involving surgery and/or whole brain radiotherapy, while effective, can significantly impair quality of life. Stereotactic radiosurgery can provide excellent control but continued development of new brain metastases remains a challenge.^2,5^ Despite molecular-targeted agents that penetrate the blood-brain barrier, disease progression in the brain remains a problem. The lack of preclinical NSCLC brain metastasis models is one barrier that may be preventing the development of effective therapeutic strategies.

Here we describe the development of a panel of NSCLC brain metastasis PDXs and demonstrate their preclinical utility in evaluating therapies. We accomplished a high engraftment efficiency, and showed that these PDXs closely resemble the patient’s original metastatic tumor by retaining their morphologic and molecular features. In addition, we demonstrated the capability of these tumors to grow as cell lines and as intracranial implants making them a valuable resource to study brain metastases. To expand upon their translational potential, we characterized the PDXs response to radiotherapy, demonstrated tumor growth inhibition with MEK inhibition in *KRAS* mutated PDXs and showed significant growth delay with the *MET* inhibitor savolitinib in a PDX harboring a MET exon 14 mutation. Furthermore, combining savolitinib with clinically meaningful doses of radiation resulted in radiosensitization in our MET exon 14 mutated NSCLC PDX. This combination provides a potential treatment strategy for patients with brain metastases from exon 14 mutated NSCLC.

PDXs have both advantages and disadvantages compared to other NSCLC brain metastasis models. PDXs allow for quick testing of molecular-targeted therapies, recapitulate human tumors and are cost effective. Other available models include human and murine cell line xenografts, spontaneous murine models, and humanized mouse xenografts.

Human cell line models generally show a homogeneous histology and lack human stroma, which is important for modeling the metastatic process. These cell lines are derived by serially passaging cells in culture, inoculating them in immunodeficient mice and selecting cells that seed the brain. There are two known available NSCLC cell lines that have tropism for the brain, H2030-BrM3 and PC9-BrM3, both established from lymph nodes of patients with lung adenocarcinoma and not from brain metastasis tumors.^38^ Syngeneic murine cell line models allow for assessment of immunotherapies but are dependent on mouse tumors cell lines, which are typically driven by specific driver mutations and may not reliably recapitulate human disease. The only reported syngeneic murine cell line model used Lewis lung carcinoma cells either directly implanted into the brain or forming brain tumors through hematogenous spread by intracardiac injections.^39^

Spontaneous brain metastases can be modeled in immunocompetent, genetically engineered mice (GEM) by manipulating oncogenes or tumor suppressor genes.^40,41^ Currently there are no available NSCLC brain metastasis GEM models; however, two models of melanoma^42,43^ and one model of small cell lung cancer^44^ have been reported. In these models, the formation of metastases is limited because primary tumors grow quickly and animals usually have to be euthanized before brain metastases develop.^45^

Compared to cell line and GEM models, PDXs are directly established from patients and thus closely reproduce the molecular, genetic and histological heterogeneity of the original tumor. The disadvantage of PDXs is the lack of an immune system – however, this can be overcome by using humanized mouse PDX models. Humanized mouse models engraft components of the human immune system into otherwise immunocompromised mice.^46^ Ideally, humanized mice would have the same immune system from which the PDX was derived. However, it is difficult to obtain CD34+ cells from cancer patients, and therefore, an allogeneic immune approach is typically used.^47^ These are complex models and currently no humanized mouse models for NSCLC brain metastasis exist.

In conclusion, we have developed a panel of human derived NSCLC brain metastasis xenografts and cell lines, which have the capability of forming intracranial tumors. These PDXs display a spectrum of differentiation typical of clinical pulmonary adenocarcinoma samples, and a high degree of phenotypic and genetic consistency between original patient tumor and xenograft. The use of early passage xenografts and cell strains provides a powerful system for investigating novel therapeutics and combination therapies, and for testing hypotheses of mechanisms of therapeutic response and resistance in human NSCLC brain metastases.

## Supporting information

Table S1: Detailed mutation list

Table S2: Short tandem repeat analysis loci

Table S3: KEGG (Kyoto Encyclopedia of Genes and Genomes) Pathways

Table S4: Gene Ontology Terms

## Acknowledgements

We would like to thank Dr. M. Shahriar Salamat and University of Wisconsin Department of Pathology and Laboratory Medicine for initial tissue processing of patient samples. We also would like to acknowledge the University of Wisconsin Carbone Cancer Center Translational Science BioCore Biobank for providing tumor samples for PDX development.

## Disclosures

none

## Funding

This project was supported in part by grants from the American Cancer Society (RSG-16-091-01-TBG), the UW Paul P. Carbone Young Investigator Award, the University of Wisconsin Lung Disease Oriented Team, and the University of Wisconsin Carbone Cancer Center Support Grant (P30 CA014520).

## Supplemental Tables

Table S1: Detailed mutation list

Table S2: Short tandem repeat analysis loci

Table S3: KEGG (Kyoto Encyclopedia of Genes and Genomes) Pathways

Table S4: Gene Ontology Terms

## Notes

### Competing Interest Statement

The authors have declared no competing interest.

### Summary of Updates

New figure 5 has been added.

