## Supplementary material for "Development and Characterization of Patient-Derived Xenografts from Non-Small Cell Lung Cancer Brain Metastases": Table S1: Detailed mutation list

**Supplemental Table 1: Detailed Mutation List**

| **PDX#** | **Histology** | **PD-L1** **expression** | **Tumor Mutation Burden** | **Mutation Profile** |
| --- | --- | --- | --- | --- |
| **UW-lung-2** | adenocarcinoma | 70% | 19 Muts/Mb | KRAS G12C  TP53 splice site 994-2A>T  CDKN2A p16INK4a E119*  p14ARF *133Lext*23  RBM10 splice site 1575+1G>C  SMARCA4 1548G>C, 2735_2736delGCinsTG  ATM 4626G>C  KEAP1 599A>C  NFKBIA amplification  NKX2-1 amplification  MYC amplification |
| **UW-lung-3** | adenocarcinoma | 20% |  | NOTCH4 1099G>T  PLCG2 3616C>T  KMT2D 10993C>G |
| **UW-lung-4** | adenocarcinoma | 15% | 25 Muts/Mb | KRAS G12V 35G>T  ATM splice variant 3285-2A>T  FBXW7 337_339delGAG  NFE2L2 101G>A  RBM10 1285C>T  SF3B1 1998G>T  GNAQ 943G>T  JAK2 1993G>C  LRP1B 10532-16_10532-15insT, 2966C>A  PIK3C2B 260C>G  KMT2D 9557G>T |
| **UW-lung-5** | adenocarcinoma | <1% | 9 Muts/Mb | KRAS G12C 34G>T  STK11 splice site 465-2A>T  CDKN2A loss  IGF1R amplification – equivocal  KEAP1 splice site 1708+1G>T |
| **UW-lung-7** | adenocarcinoma | 50% |  | TP53 379T>C, 718A>G – subclonal  DNMT3A 2478+13G>A  KMT2D 4021-15_4021-14delCT  AKT2 amplification  CCNE1 amplification |
| **UW-lung-16** | adenocarcinoma | <1% |  | KRAS G12C 34g>T  TP53 R213* 637C>T  STK11 40delG frameshift  CDKN2A 322G>A  APC 1240C>T  CHECK2 1556C>T  MTOR 3676G>C, 1198G>A  PIK3CA 733G>C  KEAP1 679C>T stop gained |
| **UW-lung-18** | adenocarcinoma | 35% |  | KRAS G13C 37G>T  TP53 C135Y 404G>A  KDR 3872G>T, 2516C>A  RAD54L 604C>T  KEAP1 1435G>T |
| **UW-lung-20** | sarcomatoid carcinoma with adenocarcinoma component | 100% | 9.2 Muts/Mb | TP53 c.376-2A>t Splice region Variant - LOF  TP53 p.R2495 Missense variant-LOF  CKS1B copy number gain  CDKN2A 206A>G  BCL6 729_731dupCAG  ARID2 3151C>T, 3337G>T  HGF 685G>T  MET Asp1000 frameshift |
| **UW-lung-21** | Adenocarcinoma | 100% |  | MET exon 14 deletion  MET exon 14 splice site mutation  TP53 1025G>C  CCND1 amp 7  CDKN2A 206A>G  TERT 228C>T |

PD-L1, programmed death-ligand 1; PDX, patient-derived xenograft; UW, University of Wisconsin
