## Supplemental Methods for "Development and Characterization of Patient-Derived Xenografts from Non-Small Cell Lung Cancer Brain Metastases"

*In vivo* immunohistochemistry

5 μm sections from formalin fixed paraffin embedded samples were deparaffinized with Xylene and hydrated through graded solutions of ethanol. Antigen retrieval was conducted in sodium citrate retrieval buffer (pH 6.0) followed by washing in running water. Slides were washed in PBS and then incubated with 0.3% hydrogen peroxide solution. Blocking was carried out using 10% goat serum in PBS and then incubated with the primary antibody (Supplemental Table) diluted in 1% goat serum in PBS containing 0.1% Triton X-100 overnight at 4°C. Slides were washed with PBS next day; secondary antibody was used (SignalStain® Boost IHC Detection Reagent (HRP, Rabbit) CST #8114). Staining was detected using diaminobenzidine (Vector Laboratories, Inc. #SK-4100). The slides were counterstained with 1:10 hematoxylin (Thermo Scientific #TA-125-MH) solution for 2 minutes, then dehydrated in ethanol and xylene solutions and sections were covered with coverslip with Cytoseal (Thermo Scientific #8312-4).

Antibodies used in this study

| **Antibody** | **Source** | **Company** | **Catalog no.** | **Dilutions** |
|---|---|---|---|---|
| TTF-1 (SPT24) | Mouse | BioCare | ACI3126C | 1:100 |
| Ki-67 (D2H10) | Rabbit | Cell Signaling | 9027 | 1:400 |
| phospho-p44/42 MAPK (Erk1/2) (Thr202/Tyr204) | Rabbit | Cell Signaling Technology | 4370 | 1:200 |
| phospho-S6 Ribosomal Protein (Ser235/236) | Rabbit | Cell Signaling Technology | 4858 | 1:400 |
| MET (D1C2) | Rabbit | Cell Signaling Technology | 8198 | 1:320 |
| Phospho-Met (Tyr1234/1235) | Rabbit | Cell Signaling Technology | 3077 | 1:320 |
